# Convergent within-host evolution alters key virulence factors in a *Klebsiella pneumoniae* clone during a large hospital outbreak

**DOI:** 10.1101/2024.02.06.577356

**Authors:** Greta Zaborskytė, Karin Hjort, Birgitta Lytsy, Linus Sandegren

**Affiliations:** Department of Medical Biochemistry and Microbiology, Uppsala University, Uppsala, Sweden.; Department of Medical Sciences, Uppsala University, Uppsala, Sweden. Current address: Department of Laboratory Medicine, Division of Clinical Microbiology, Karolinska Institute, Solna, Sweden

## Abstract

Bacterial pathogens adapt to host niches because of within-host selective pressures, and this evolutionary process provides valuable insights into host-pathogen interactions. However, genetic changes underlying adaptive phenotypes are difficult to identify from data generated by genome-wide association studies of unrelated bacterial clones. Here, we followed the evolution of a single *Klebsiella pneumoniae* clone in 110 patients during a 5-year nosocomial outbreak by combining comparative genomics with phenotypic characterization. Strong positive within-patient selection targeted key virulence factors in isolates from infection sites. The clone repeatedly lost acute virulence primarily via alterations in capsule and lipopolysaccharide, changed regulation of iron uptake, and increased biofilm formation. These phenotypes represent likely niche adaptations, mainly to the urinary tract, and some were associated with trade-offs during gastrointestinal colonization. The substantial convergent evolution reflects the trajectories undertaken by high-risk clones of *K. pneumoniae* and other pathogens adapting during acute and chronic infections.

## Introduction

Opportunistic pathogens like *Klebsiella pneumoniae* often colonize the gastrointestinal tract (GI) or nasal mucosa without causing disease^1,2^. However, different tissue compartments (e.g., the urinary tract, lungs, blood) can be invaded due to immune suppression, indwelling medical devices, and antibiotic treatment^3,4^. During colonization and infection, bacterial populations encounter numerous challenges, such as obtaining essential nutrients like iron, need for binding to cells or abiotic surfaces, and surviving attacks from the immune system. This creates a potential for within-host selection of mutant subpopulations that are better fit for particular host niches^5–11^. The fate of such mutants depends on their survivability in the new niche and potential trade-offs to their normal colonization and transmission routes. While ”adapt-and-live” mutations maintain pathogen transmission and are associated with globally disseminated clones, “adapt-and-die” changes promote survival in a specific host niche but limit bacterial spread to a new host due to trade-offs or growth in a physical location from which spread is limited^12,13^. Evolutionary studies on *K. pneumoniae* mainly focus on horizontally transmissible genetic traits that lead to the emergence of high-risk clones, whereas data on transient adaptations selected on the individual host level are scarce. However, such transient adaptive changes may affect the outcome of an infection in the individual patient, for example, by prolonging it or making it more severe^9–11^.

Often, multiple diverse clones of *K. pneumoniae* circulate in the same hospital and infect patients simultaneously^14,15^. While large-scale genomic comparisons of such unrelated bacterial isolates can be useful, differentiation between background allelic variation and true adaptive changes becomes complicated, thus reducing the resolution of the within-host evolutionary trajectories. In contrast, clonal hospital outbreaks allow for a more precise analysis due to the common genetic origin of bacteria isolated from different patients and the defined time frame^16,17^.

During 2005-2010, an outbreak of a multiresistant *K. pneumoniae* clone occurred at Uppsala University Hospital, with 320 identified patients either colonized or infected^18–20^. Transmission of this outbreak clone occurred via direct person-to-person contact and not from contamination of hospital surfaces^19^; therefore, genetic changes under positive selection observed among isolates should illustrate adaptive evolution to the host rather than to the environment. Here, we combined large-scale genomic and phenotypic analysis to understand the within-host evolutionary trajectories of the *K. pneumoniae* clone in 110 patients during asymptomatic colonization and infection. We find strong positive selection acting on key virulence factors especially in isolates from infections and show how the genetic changes translate into diverse adaptive phenotypes connected to survival in the host.

## Results

### Strong positive within-patient selection targets key virulence factors at infection sites

We determined the complete genome sequence of the index isolate and 109 representative infecting and colonizing isolates distributed throughout the outbreak (Fig. 1a and Supplementary Fig. 1). Single nucleotide polymorphisms, insertions, and deletions were present in 407 chromosomal genes and 91 intergenic regions with a total of 653 unique genetic changes equalling an average rate of 3.0 mutations per genome per year (Fig. 1b and Supplementary Tables 1 and 2). When analyzing infecting and colonizing isolates separately, the infection isolates showed a 3-fold higher mutation rate. Phylogenetic analysis generated a tree topology dominated by individual branches and a few clusters (A-H), each with long individual branches, indicative of selection of genetic changes within individual patients rather than spread of dominating bacterial lineages (Fig. 1c). The ratio of nonsynonymous vs. synonymous changes in protein coding genes (dN/dS) showed a clear positive selection signal with an overall dN/dS of 5.7 and a very strong signal (dN/dS = 93) for the 19 genes with three or more independent mutations. There was a strong overrepresentation of nonsynonymous mutations in the branch “tips” rather than in clusters, supporting independent acquisition of adaptative mutations among isolates (Supplementary Fig. 2).

**Fig. 1.**
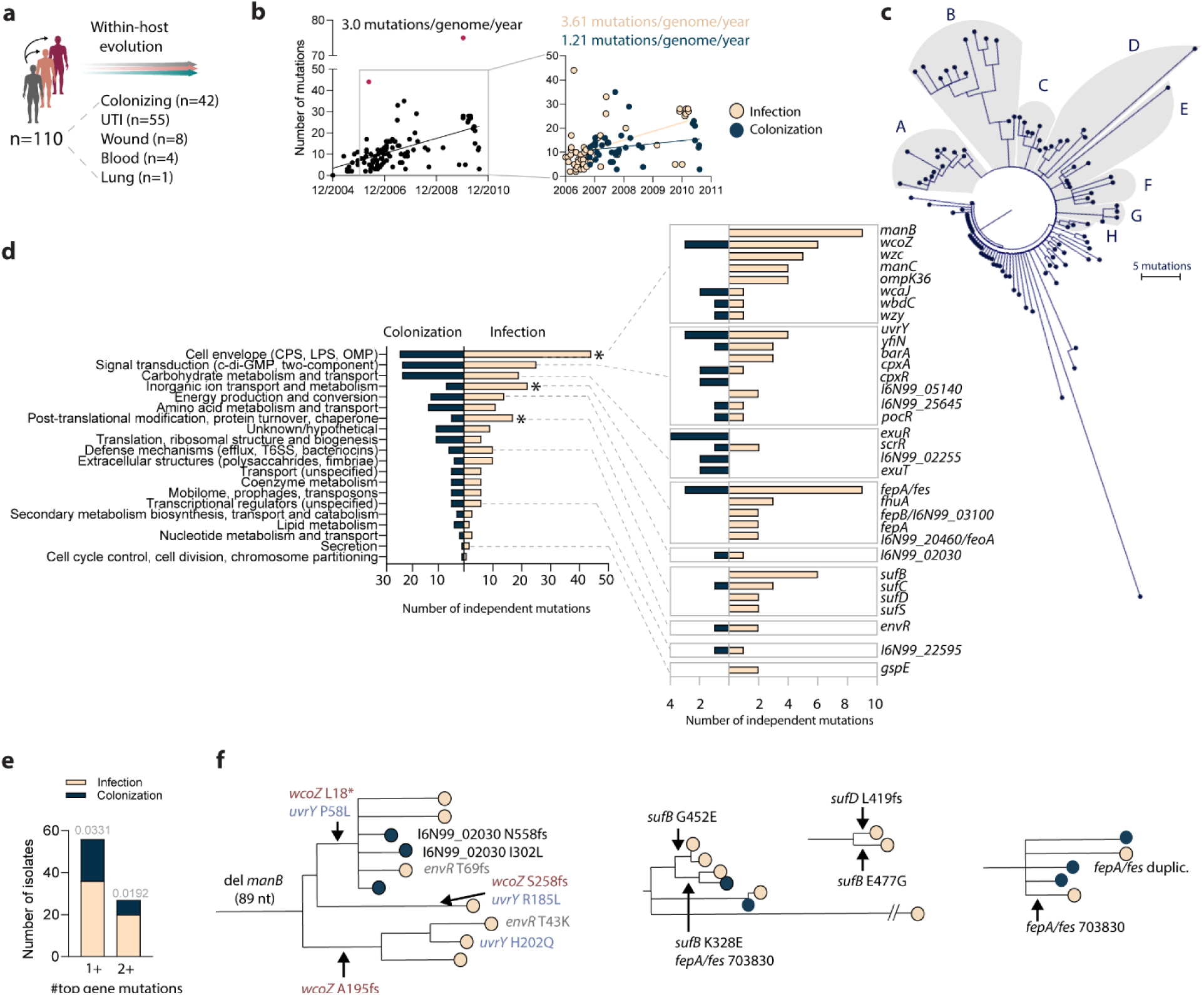
| Strong positive selection of genetic changes during infection. **a,** Overview of isolates included in the study, **b,** Genetic changes per isolate relative to the index isolate plotted over time of isolation. Red isolates (DA25065 and DA69557) were excluded from the calculation of mutations/ge-nome/year because the number of genetic changes exceeds the mean plus two standard deviations. The insert shows infection and colonization isolates separately during the period when both types of isolates were collected {starting in September 2006). **c,** Maximum likelihood phylogeny of outbreak isolates. Clusters of isolates are shaded and assigned to letters A-H. **d,** Functional grouping of genes with at least two nonsynonymous mutations selected independently as tip mutations. Asterisks denote groups with a significant bias between infecting and colonizing isolates (binomial test, *p<0.05).The intergenic regions not assigned to any of the functional groups can be found in Supplementary Table 3. **e,** Number of colonizing and infection site isolates with at least one (1 +) and 2 or more (2+) top genes and intergenic regions mutated. Bias between infection and colonization was tested using the binomi­al test, p values are depicted above the columns, **f,** Examples of convergent mutation selection within phylogenetic clusters.

The majority of infecting isolates were from urinary tract infections (UTIs). While rich sugar-based carbon sources and competition with other microbiota prevail in the intestines, the urinary tract is typically a nutrient-poor environment dominated by amino acids, limited iron, and fierce host defences^21^. Therefore, the adaptations selected in the gut versus urinary tract would likely display a functional bias. Independently selected genetic changes in cell envelope, iron uptake and metabolism, and Fe-S cluster synthesis were significantly overrepresented among infection isolates (Fig. 1d). This bias was even more pronounced on an individual gene level when considering genes and intergenic regions (n=39) with at least two mutations acquired in parallel in different isolates (two-tailed binomial test, p<0.0001). Adaptation to specific niches often requires multiple mutations that produce a combination of altered traits^22^. Here, we find that genetic changes in the most frequently mutated genes often co-occurred with up to eight top genetic targets mutated in the same isolate (Fig. 1e, Supplementary Fig. 3 and Supplementary Table 3). This co-occurrence was also overrepresented in infection isolates, mainly from UTIs, further supporting strong convergent niche-specific evolution. Some convergent mutations were also acquired at different branching points within the same phylogenetic cluster, e.g., *wcoZ* (capsule acetyltransferase), *uvrY* (transcriptional regulator in UvrY-BarA), *envR* (efflux pump regulator), *suf* (Fe-S cluster synthesis), and intergenic *fepA/fes* (siderophore uptake) (Fig. 1f). Collectively, comparative genomics shows that changes in key virulence-associated processes were repeatedly selected in individual patients in parallel and preferentially during infection.

### Frequent attenuation in acute virulence and increased susceptibility to complement due to repeatedly selected surface polysaccharide mutations

Considering the strong positive selection acting on vital virulence factors in infecting isolates, we screened all 110 isolates for their potential to kill *Galleria mellonella* wax moth larvae. *G. mellonella* has an innate immune system with both cellular and humoral responses similar to those of vertebrates^23,24^ and is the preferred infection model for opportunistic *K. pneumoniae,* which do not perform well in mice models^25–27^. Based on the rate of *G. mellonella* killing and the health status of the surviving larvae, the isolates were divided into five groups (Fig. 2a). Half of the isolates showed complete or partial attenuation, while an increased virulence was seen for five isolates not associated phylogenetically. One group of isolates (phylogenetic cluster B) displayed a different infection progression, where injected larvae showed no signs of sickness up to 12 h post-injection but died rapidly within 24 h. The attenuated group was overrepresented among infection site isolates (Fig. 2b), strongly indicating a switch to a more chronic form of infection as a common adaptation.

**Fig. 2.**
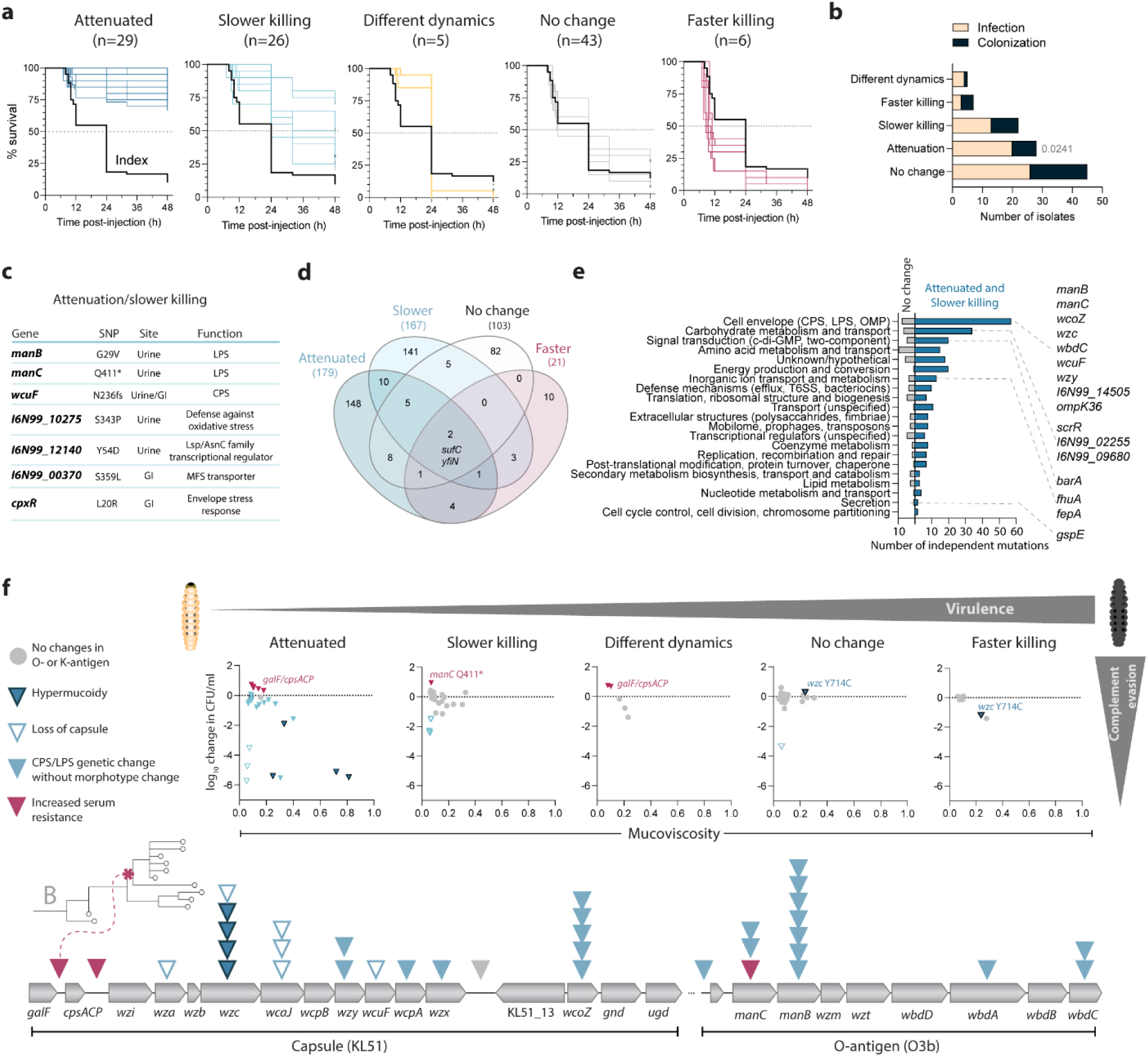
| Virulence profiles show frequent attenuation and occasional increase. **a**, Kaplan-Meier survival curves of G. *mellonel/a* larvae infected with bacterial isolates (n=20 larvae for each isolate). Assignment of virulence groups is based on larval survival and the cumulative health index scores of the surviving larvae after 48 h, compared with the index isolate (log-rank Mantel-Cox test), **b,** Distribution of isolates from infection and colonization among the virulence groups (binomial test), **c,** Single mutations identified as responsible for acute virulence loss early stop codon, “fs” - frameshift). **d,** Venn diagram showing the overlap among the genetic targets (genes and intergenic regions) between isolates with different virulence profiles. Each genetic change was categorized as possibly contributing to the virulence based on the isolate’s virulence profile and its phylogenetic location, **e.** Functional groups of unique mutations in the “no change” and “atΐenuation7“slower killing” groups. Named genes acquired at least two independent mutations in attenuated/slower killing isolates, **f,** Upper panel: bacterial survival in human serum (loglO change in CFU/ml after 3 h) versus sedimenta­tion (mucoviscosity) in different virulence groups. Lower panel: independent mutations in cpsand O-antigen regions. Part of phylogenetic cluster B has an intergenic mutation *galF/cpsACP* conferring resistance to serum.

In some cases, it was easy to pinpoint the single mutations responsible for acute virulence loss by comparison of neighbors in phylogenetic clusters (Fig. 2c). For most isolates, though, there was more than one genetic difference between isolates with different virulence profiles, complicating the distinction between virulence-altering changes and co-selection. There was minimal overlap between genes and intergenic regions mutated in different virulence groups, with most unique hits in the isolates with impaired acute virulence (Fig. 2d). These unique mutations were associated with diverse functions, but parallel cell envelope mutations were the dominant ones, particularly in surface polysaccharide synthesis regions (*cps* and O-antigen) (Fig. 2e, f and Supplementary Table 4). Often, combinations of the same mutational targets were acquired in single isolates, even with identical combinations (e.g., *wzc*+*manB*+*ompK36*). No unifying mutations were present in the increased virulence group, reflecting diverging rather than converging genotypes associated with a gain of function (Supplementary Table 4).

Capsule typically shields *K. pneumoniae* from complement attacks, which is an essential innate immune response to protect against infection^28^. Therefore, a non-mucoid morphotype (capsule loss) has been associated with impaired complement evasion, while increased mucoviscosity has been linked to complement resistance and more invasive infections^29^. Surprisingly, here the most hypermucoid isolates were attenuated and as sensitive to human serum as the isolates that had lost the capsule, with up to >5 log reduction in CFU/ml within 3h (Fig. 2f). All such hypermucoid isolates carried missense mutations in *wzc,* encoding an essential capsule co-polymerase and regulator^30,31^. The exception was two clustering isolates with the *wzc*^Y714C^ mutation. Increased mucoviscosity due to other capsule changes, e.g., a 21-nt deletion in the capsule regulator *yrfF*, did not reduce resistance to complement. All *wzc* mutants were isolated from infection sites, suggesting a unique adaptation.

In most cases, reduced killing of *G. mellonella* was coupled with decreased complement evasion (Fig. 2f). However, increased complement resistance via the O-antigen mutation *manC*^Q411*^ and an intergenic capsule mutation associated with *galF/cpsACP* also resulted in slower killing of *G. mellonella*. This finding suggests that certain modifications of surface polysaccharides could help avoid detection, thereby delaying the host response. Within some phylogenetic clusters, sensitivity to complement via capsule loss was acquired subsequent to attenuation in *G. mellonella* in colonizing isolates (Supplementary Fig. 4). Moreover, some attenuated isolates had non-altered survival in serum, which suggests other factors than different interaction with complement as the basis for virulence loss. One such contributing factor could be a general growth defect^25^, although we did not see any correlation between killing of *G. mellonella* and the exponential growth rates *in vitro* (Supplementary Fig. 5).

### Intergenic *fepA/fes* changes alter the regulation of enterobactin uptake and utilization

Iron sequestration is essential for the growth and survival of *K. pneumoniae* inside the host^32–34^. Multiple siderophore-dependent and independent systems are responsible for this process (Fig. 3a), and all of them were mutational targets among the isolates, e.g., inactivation of *fhuA*, nonsynonymous mutations in *fepA*, gene amplifications of *feoABC*, and intergenic changes in the *fepA/fes* regulon (Supplementary Table 2). Moreover, they were often co-selected with mutations in *suf* genes, especially *sufB* and *sufC*, responsible for Fe-S cluster synthesis (Supplementary Fig. 3). We measured how well all isolates grew under iron-depleted and repleted conditions with either FeSO_4_ or inactivated human serum as a source of iron. While the overall growth rates and yields were decreased in the absence of iron, we did not observe any apparent connection between mutations and changes in iron-dependent growth (Supplementary Table 5). Therefore, seemingly loss-of-function changes did not result in defects during iron-dependent growth, possibly because partially redundant systems complement measurable deficits.

**Fig. 3.**
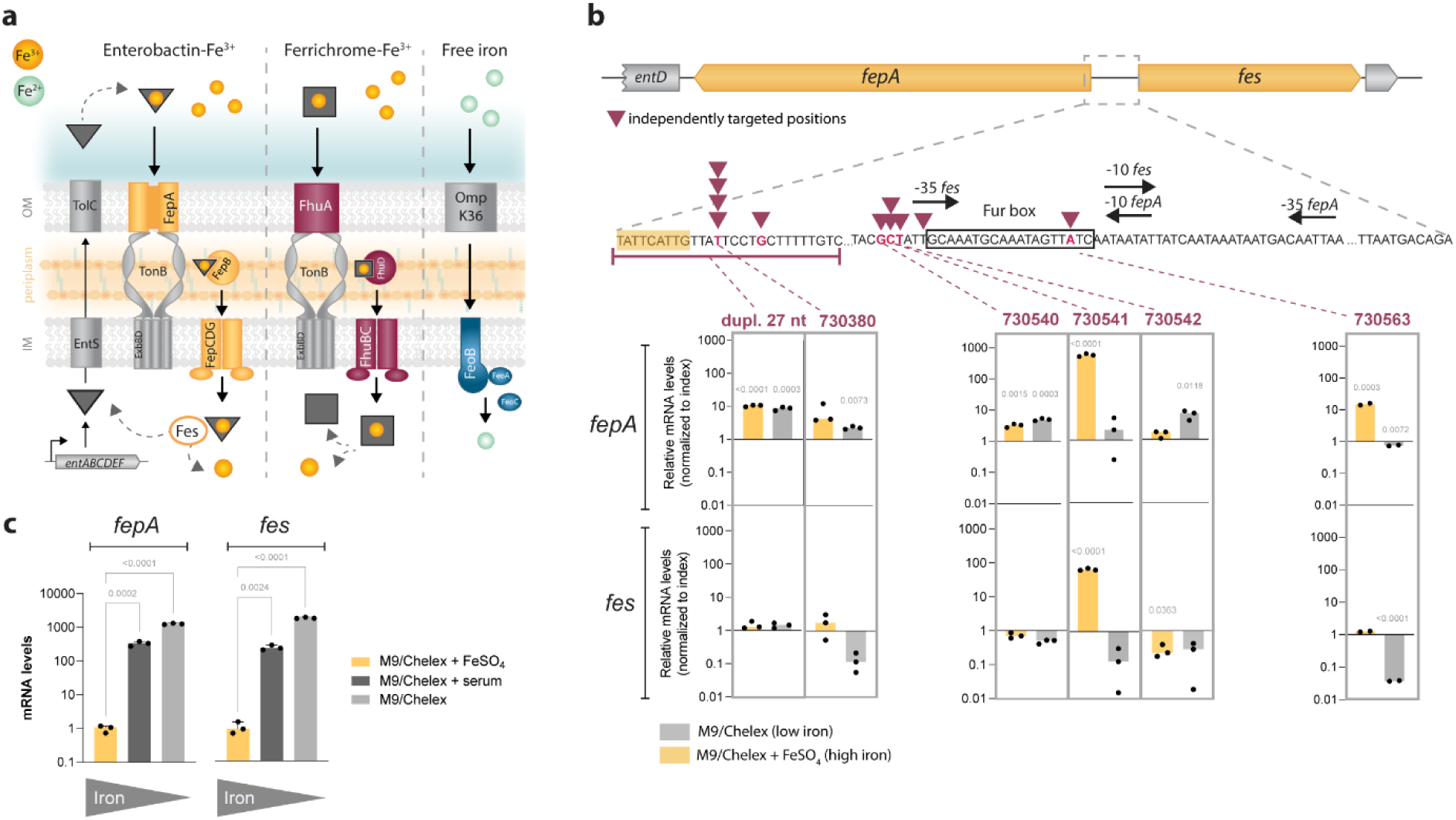
| Regulatory changes in iron uptake and utilization. **a**, Schematic overview of *K. pneumoniae* iron uptake systems for iron bound to sidero-phores, such ascatecholate-typeenterobactin or hydroxamate-typeferrichrome, and free ferrous iron, **b,** Regulatory elements in the *fepA/fes* region and changes in *fepA* and *fes* mRNA levels in isolates with mutations (noted in dark red) relative to the index isolate. Comparison to the index isolate in each condition was done by unpaired t-test; significance at p<0.01 after Bonferroni correction for five comparisons. **c,** Changes in *fepA* and *fes* mRNA levels in the index isolate under low and high iron conditions. Results are shown for three biological replicates. mRNA levels in **b** and **c** are relative to the house­keeping gene *glnA* and further normalized to M9/Chelex + FeSO_4_. Comparison between low and high iron conditions was done by one-way ANOVA followed by Dunnett’s multiple comparison test; p values are depicted above the columns.

The *fepA/fes* region was the most common independently targeted intergenic region (Fig. 1d, f), but also intergenic regions upstream of *fepB* and *feoA* were mutated, mostly in infecting isolates. The *fepA/fes* intergenic region contains overlapping bidirectional promoters and a Fur (ferric uptake regulator) box that regulates the expression of the FepA enterobactin transporter and the Fes enterobactin esterase (Fig. 3b). As expected, the index isolate regulated the expression of both genes depending on iron levels with high expression in chelated M9 medium, low expression in chelated M9 supplemented with FeSO_4_, and an in-between expression in medium supplemented with serum where most iron is bound to proteins (Fig. 3c). Different mutations in the region right upstream of *fepA* or around and inside the Fur box either increased *fepA* expression or eliminated the responsiveness to iron availability such that the expression of both *fepA* and *fes* remained elevated even in high-iron conditions, most likely through disruption of the Fur-mediated regulation (Fig. 3b). Taken together, these results illustrate that highly specific phenotypic responses mediated by intergenic mutations associated with iron acquisition were selected during infection.

### Increase in biofilm formation on catheter-like surfaces were selected via mutations affecting c-di-GMP turnover, fimbriae, and polysaccharide synthesis

Most of the infection isolates were from UTIs, where urinary catheterization promotes biofilm growth of uropathogens, including *K. pneumoniae*^35–37^. Therefore, we tested how well the isolates formed biofilms on silicone-coated surfaces with and without fibrinogen using the FlexiPeg device^38^. Fibrinogen is commonly deposited on urinary catheters during catheter-induced inflammation^39^ and is required for attachment by several uropathogens^40–42^. Thirteen isolates (nine urine/wound) had elevated levels of biofilm formation and were distributed throughout the phylogeny; seven of them needed fibrinogen coating to display the phenotype fully (Fig. 4a).

**Fig. 4.**
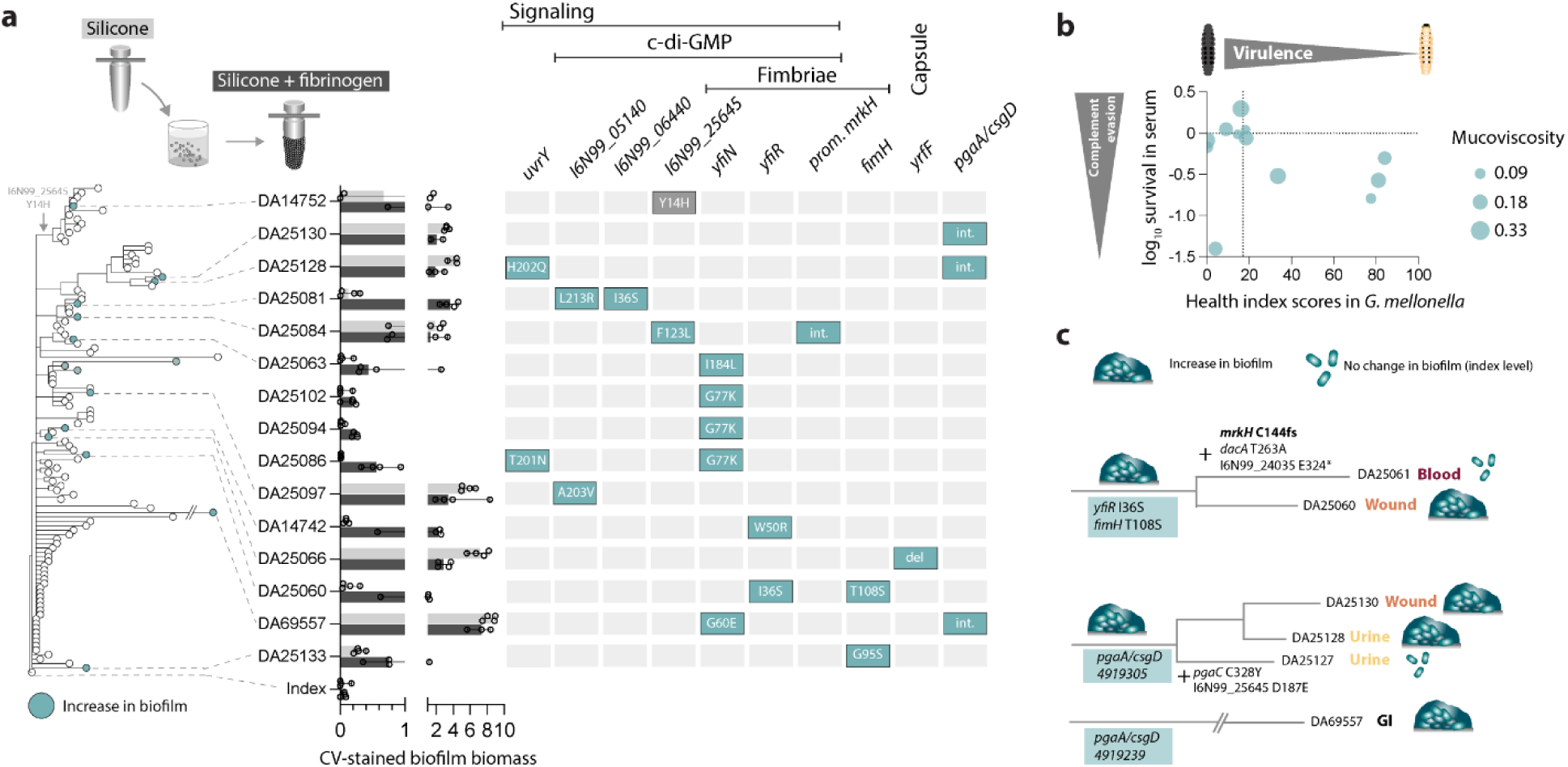
| Convergent evolution of biofilm formation capacity. **a**, Phylogenetic distribution of isolates with increased surface-attached biofīlm capacity relative to the index isolate. Bars indicate medians of total CV-stained biomass of 48 h biofilms grown on silicone-coated pegs without (light grey) or with fibrinogen coating (dark grey). At least four biological replicates (dots) are shown. Highlighted genetic changes are most likely responsible for the increased biofilm phenotype, b, Interconnection between acute virulence and survival in serum among isolates with increased biofilm formation capacity. ¢, Biofilm evolution scenarios with consecutive genetic changes illustrated on branches where mutations leading to increased (DA25060, DA25130/-DA25128, DA69557) or decreased (DA25061, DA25Ί27) biofilm formation are shown.

The mutations responsible for an increased biofilm formation were in genes affecting fimbrial adhesins, c-di-GMP signaling proteins, and polysaccharides, including capsule regulation (Fig. 4a). Adaptive changes in the FimH tip adhesin were previously reported in *Escherichia coli* and *K. pneumoniae* isolates from UTIs^43–46^, while YfiN and YfiR were targeted in *Pseudomonas aeruginosa* adapting during chronic lung infections^47^. The wound isolate DA25066 with extremely high biofilm biomass had a unique 21 nt in-frame deletion in *yrfF* that encodes a negative regulator for the Rcs two-component system, which induces capsule production^48^. Previously, a 12-nt deletion in *yrfF* has been reported in *E. coli* isolated from keratitis^49^ and selected during serial passaging in macrophages^50^. The isolate DA69557 had multiple mutations with possible connection to an extreme biofilm capacity previously observed by SEM^38^. One such change was the intergenic *pgaA/csgD* mutation (poly-N-acetylglucosamine PNAG exporter/master biofilm regulator) that was acquired in parallel both in DA69557 and part of cluster B. Moreover, at least four different genes associated with c-di-GMP turnover, including *yfiN*, were mutated (Supplementary Table 1).

There was no direct relationship between acute virulence features (*G. mellonella*, serum, mucoviscosity) and biofilm capacity in these isolates (Fig. 4b). In most cases, the increased biofilm capacity had evolved as the end-point adaptation; however, two clusters showed reversion of this phenotype (Fig. 4c). The blood isolate DA25061, found 1.5 months later than the phylogenetically closely related wound isolate DA25060, lost the increased biofilm phenotype due to an inactivating mutation in the transcriptional activator MrkH. Similarly, a mutation in the essential PNAG synthesis protein PgaC reverted the biofilm phenotype back to the index level in part of cluster B that otherwise showed an increase due to the intergenic *pgaA/csgD* mutation. Such observations reflect likely differences in biofilm-related fitness at different body sites (e.g., wound vs. bloodstream), but also how individual host environments shape bacterial evolution even in the same niche (e.g., urinary tract). Differences in the immune response have been suggested to affect the adaptive trajectories during asymptomatic bladder colonization with *E. coli*^6^.

### Increased biofilm capacity gives an advantage during antibiotic-facilitated GI colonization in mice

Since intestinal colonization is usually the dominant reservoir of spread to new hosts and also precedes infection by classical *K. pneumoniae* strains^2,51,52^, we tested how a set of isolates with different evolved phenotypes compete with the index isolate when colonizing the GI tract of mice. Colonization with the natural gut microflora present was not stable in any of the competition groups, indicating that, at least under these conditions in mice, the outbreak clone is not a super colonizer (Fig. 5a). When the drinking water was supplemented with clindamycin, which causes significant disruptions to the gut microbiota and enables long-term colonization with *K. pneumoniae*^53^, the bacterial load dramatically increased above 10^9^ CFU/g faeces and stayed so until the end of the experiment. These results signify the role of antibiotic treatment-induced alterations to the gut microbiota in colonization with the outbreak clone, supporting its efficient spread in the hospital.

**Fig. 5.**
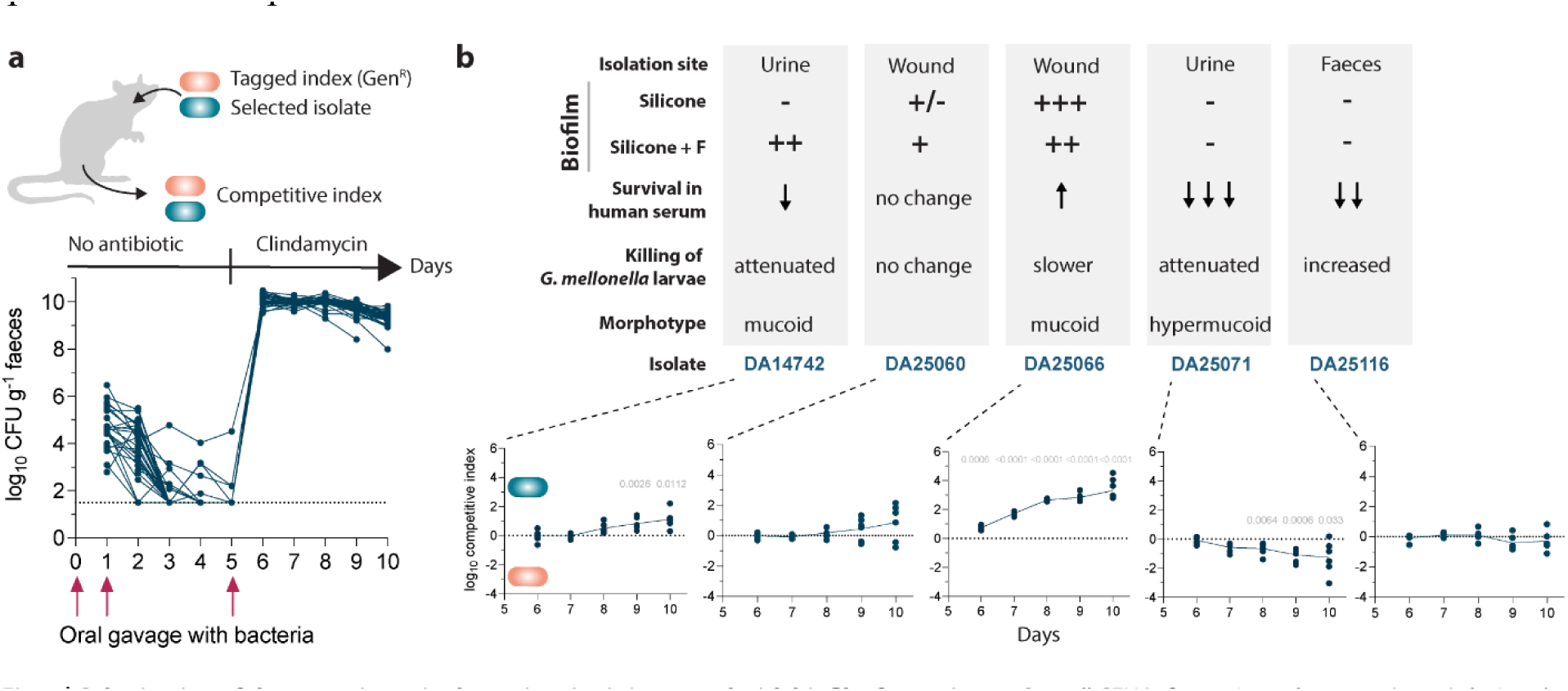
| Colonization of the gastrointestinal tract in mice is increased with biofilm formation. **a**, Overall CFU/g faeces in each mouse (n=35) during the 10-day experiment. On Day 5, clindamycin was added to the drinking water after a third oral gavage with bacteria, **b,** Top panel illustrates phenotypes for each isolate. Lower panel displays competitive indices during days 6-10 of gastrointestinal colonization. Competitive index >0 means an advantage of the tested isolate versus the index isolate DAΊ4734, <0 - disadvantage. Dots show the competitive index in individual mice (n=6 per competition) and the trendline connects the means. Comparisons to control competition on each day were made by unpaired two-tailed t-test; significance after Bonferroni correction for multiple comparisons at p<O.OΊ, p values depicted above the datapoints.

All three biofilm-forming isolates had a selective advantage over the index isolate during GI colonization. This was especially prominent for DA25066, with the most substantial biofilm formation capacity conferred by a deletion in the capsule regulator *yrfF* (Fig. 5b). Two of the biofilm-forming isolates also carried a mutation in *sbmA*, encoding the uptake protein for microcins, suggesting possible alterations in the defense against other gut bacteria. The colonizing isolate DA25116 stayed neutral, and the hypermucoid *wzc* mutant DA25071 was not as good at colonizing GI as the index isolate. Therefore, specific phenotypes (e.g., biofilm formation) selected during within-host evolution favor GI colonization, while others (e.g., *wzc*-mediated hypermucoidy) do not. This observation reflects how, given the opportunity, mutants with an advantage at infection sites could be pre-selected already during gut colonization and have an advantage in both body sites. In contrast, a hypermucoid *wzc* mutant isolated from a UTI represents a case where the mutant had adapted to the infection site so well that it no longer can colonize the gut to the same extent (“adapt and die”).

## Discussion

Understanding within-host evolutionary dynamics of bacterial populations is central to infection biology. Our study reconstructs key genomic and phenotypic events in 110 isolates of a single *K. pneumoniae* clone during transmission between patients and adaptation to host niches. The adaptive changes we identify as selected mainly in infecting isolates overlap with those linked to high-risk *K. pneumoniae* clones^54–56^, uropathogenic *E. coli*^57^, and chronic infections by CF lung pathogens, like *Burkholderia dolosa*^58^, *B. cenocepacia*^59^, and *P. aeruginosa*^47^. While genomic analysis alone can be a powerful predictor of adaptive evolution, we show unexpected coupling to phenotypes (e.g., *wzc*, *manB/manC*, *fhuA*) in quite a few cases. These findings highlight the importance of connecting genotype-phenotype when assessing complex selection in the host^60,61^ even in a single clone.

We show that reduced evasion of innate immunity mechanisms (complement) and attenuation in acute virulence was a much more common evolutionary direction than increased virulence. Most attenuated isolates were from infection sites, suggesting that they are still maintained rather than cleared, possibly by avoiding detection by the host. Moreover, acquisition of parallel mutations and similar mutation-combinations strongly implies convergent adaptation to a more chronic state, like for other opportunistic pathogens^62^. How exactly the selected mutations mediate such a switch mechanistically in *K. pneumoniae* is a key question for future research. The assessment of increased killing of *G. mellonella* is more difficult as the infection progresses fast in the index isolate even at a relatively low infectious dose; therefore, there is a possibility that we underestimate the potential for increased virulence.

Virulence changes were often connected to capsular changes. One of the most striking cases was that *wzc*-mediated hypermucoviscosity sensitized the clone to complement and attenuated rather than increased virulence. In contrast, a recent study associated *wzc* mutants in the *K. pneumoniae* ST258 global high-risk lineage with increased macrophage resistance and higher virulence in zebrafish larvae and murine UTIs^54^. The same authors also suggested that such mutations are associated with blood infections, while capsule-loss mutants are associated with UTIs. However, here we detect both morphotypes in UTIs, which could reflect differences in strain background. Supporting the connection to UTIs, another recent study showed how urine modulates the capsule and maintains the hypermucoid phenotype of *K. pneumoniae wzc* mutants^63^.

Many seemingly loss-of-function LPS and capsule mutations altered virulence without changing the morphotype. The most prominent case was frequent frameshifts and deletions in *manB/C,* encoding the ManB phosphomannomutase and the ManC mannose-1-phosphate guanylyltransferase. Inactivation of *manB/C* abolishes capsule production in other *K. pneumoniae* strains^64^. Since KL51 capsule lacks mannose, in this clone, the mutations in *manB/C* probably affect the synthesis of O3b antigen that contains three mannose residues^65^. Loss of O-antigen due to its immunogenicity has been reported during late stages of chronic CF lung infections^66^.

Inactivating *manC* mutations have specifically been connected to adaptations in *B. dolosa* and experimentally biofilm-evolved *B. cenocepacia*^59^. Similarly, acetylation of the capsule in *K. pneumoniae* has been associated not only with a change in immunogenicity, but also altered adhesion to surfaces^67^, and here, inactivating *wcoZ* (capsule acetyltransferase) mutations were a common cause of virulence loss. Collectively, *manB/C* and *wcoZ* mutations could reflect selection connected to the catheterized bladder environment.

Iron scavenging has been identified as a selective trait in bacterial pathogens, such as *B. dolosa*^59^ and *E. coli*^68^. In accordance with our observations, adaptive changes in siderophore uptake rather than synthesis seem to be more critical in the context of infection^69^. Changes in iron uptake in infecting site isolates were often co-selected with mutations in *suf* genes, encoding components for the Fe-S cluster assembly complex, induced under oxidative stress and iron starvation^70^. Genetic variation in *suf* genes was also found to be positively selected in the global *K. pneumoniae* outbreak clones ST258^55^ and ST147^56^, and *sufB* mutations were connected with increased iron binding^55^. Notably, mutations that could be predicted to have adverse effects on iron acquisition did not reduce growth in low-iron conditions. This may reflect redundancy in how *K. pneumoniae* obtain and use iron, as shown previously for ST258, where mutation in the conserved *entS* (enterobactin export) led to higher expression of an alternative iron transport system (hemin transport)^33,34^. Alternatively, seemingly detrimental loss in enterobactin synthesis could also be attributed to the evasion of lipocalin 2, a major innate immunity protein that specifically binds enterobactin^34^. FhuA and FepA are also receptors for phages and bacteriocins^71,72^; therefore, loss of these proteins could be selected as protection from these factors. In agreement with this hypothesis, one of the isolates with a *fhuA* mutation also had an inactivating mutation in *sbmA*, encoding the transporter for microcin-type bacteriocins.

Although the dN/dS ratio is typically considered a strong indicator of positive selection^73^, our results illustrate that intergenic changes should not be neglected. We find that the intergenic region *fepA/fes* is a prevalent mutational target, especially among UTI isolates, and is associated with significant alterations in the regulation of enterobactin uptake and responsiveness to iron levels. Intergenic regions related to iron acquisition were also found to be under positive selection in ST410 carbapenem-resistant *E. coli*^68^. A study on the adaptation of *P. aeruginosa* showed that parallel intergenic evolution predominantly targeted promoters and suggested that intergenic mutations facilitate the evolution of essential genes by minimizing systemic detrimental effects^74^. In conclusion, we show at an unprecedented level how a clone, representing a critically important nosocomial pathogen, underwent extensive parallel evolution during a large hospital outbreak. We highlight the power of combining large-scale genomic analysis with phenotyping and hope that this dataset can continue to serve as a valuable resource to expand our understanding of the evolutionary within-host trajectories of *K. pneumoniae*.

## Materials and Methods

### Bacterial isolates and sampling

The first patient confirmed with the outbreak clone was identified in December 2004 and had been treated abroad (Tehran, Iran) before being admitted to the oncology ward at Uppsala University Hospital^18^. The isolates from the outbreak were collected from routine samples at Uppsala University Hospital. Isolates (n=110, each from a different patient and from the first occasion it was isolated) were randomly included in this study to cover the entire outbreak duration and to represent the clinical and screening isolates equally. All isolates were included at times with low incidence, and a proportion of isolates were randomly chosen from time points with high incidence. In total, 320 patients were identified with the clone, but unfortunately, all isolates except the ones gathered for this study were lost due to a freezer accident at the hospital, making it impossible to include additional isolates in the study in retrospect. 45% (n=50) of the patients were male and 65% (n=60) were female. The median age was 80 years (range 23–100). Clinical isolates (n=68) were collected from urine (n=55), wounds (n=8), blood (n=4), and lungs (n=1). Colonizing isolates (n=42) were collected from fecal samples as part of the screening for carriage of the outbreak clone. The summary of all isolates and associated genomic and experimental data is available as an interactive Microreact project (https://microreact.org/project/6bg86f2tgddpz4q5xavmgh).

### Whole-genome sequencing of outbreak isolates

The index isolate DA14734 was sequenced with long-read whole genome sequencing using the Pacific Biosciences platform (PacBio, Menlo Park, CA). DNA was prepared from overnight cultures using the Qiagen Genomic-Tip 100, according to the manufacturer’s instructions. PacBio sequencing was performed at Uppsala Science for Life Laboratory on an RSII system with one SMRT cell per genome. In addition, the DA14734 genome was also sequenced with short-reads using MiSeq (Illumina, San Diego, CA) with two times 300-bp paired-end read lengths using the Nextera XT DNA library preparation kit (Illumina, San Diego, CA), according to the manufacturer’s instructions. From the index isolate, a reference genome sequence was assembled with a combination of the PacBio reads and MiSeq reads. Assembly was performed with the CLC Genomics workbench v.20 (CLC Bio, Qiagen), including the Microbial Genomics Pro Suite module. The additional 109 isolates from the outbreak were also whole-genome sequenced with the Illumina technology, and the sequences were compared to the reference genome by reference assembly using the CLC Genomics Workbench. Variant detection was performed using CLC Genomics in-built detection modules for single nucleotide polymorphisms (SNPs), insertions/deletions, and structural variations. Variations in the chromosomal sequences were used to construct a Maximum likelihood phylogenetic tree using the CLC Microbial Genomics Module Epidemiology function. An initial Maximum Likelihood Phylogeny was created with a Neighbor Joining General time reversible nucleotide substitution model. This initial tree was then manually updated based on the exact genetic changes for each isolate with branch lengths adjusted to the exact number of genetic changes compared to the index isolate. All sequences have been deposited at NCBI under bio project number PRJNA857654.

### Bioinformatic analysis of genetic changes

The mutational rate throughout the outbreak was calculated by linear regression of the number of mutations per isolate compared to the index isolate and the respective isolate’s isolation date. The dN/dS ratio was calculated for groups of mutated genes with the same number of independent mutations by dividing the total number of non-synonymous changes by the total number of synonymous changes. Identification of biological functions and grouping of genes was done by a combination of manual BLASTp^75^, STRING^76^, and EggNog 5.0^77^ analysis combined with literature searches. The genetic basis for phenotypic changes was determined by first classifying all mutations as cluster or tip mutations, and then comparing tip mutations in clustering isolates with phenotypic differences. In cases with more than one possible mutation, we focused on repeatedly mutated genes in isolates with the same phenotype.

### Exponential growth rate measurements in MH II and urine

Overnight (O/N) cultures were diluted 1000-fold in MH II broth, and 300 μl were transferred to 100-well honeycomb plates. Five biological replicates were used for each isolate. The plates were incubated in a BioScreen C (Oy Growth Curves Ad Ltd) for 16 h at 37°C with shaking with OD_600_ measurements taken every 4 min. For experiments with urine, morning urine was collected from a healthy female volunteer on six different days. The urine was centrifuged (3800xg, 10 min), sterile filtered (0.2 μm pore size), and aliquots were frozen at -20°C. For experiments, frozen aliquots from different days were thawed, centrifuged again (3800xg, 10 min), and pooled together. Exponential growth rates were analyzed using the R script-based tool BAT 2.0^78^.

### Biofilm formation

The capacity to form surface-attached biofilms was determined on the FlexiPeg biofilm device^38^ with silicone-coated pegs. To further coat the pegs with fibrinogen, the autoclaved peg lid was incubated in a flat-bottom 96-well plate with 200 µl of 100 µg/ml fibrinogen from human plasma (product number F3879, Sigma-Aldrich, in 0.9% NaCl or PBS) at 4°C overnight. Approximately 10^5^ CFU in a total volume of 150 µl were added to a 96-well plate, where the autoclaved peg lid was inserted, and biofilms were allowed to form statically in BHI for 48 h at 37°C with a medium change after 24 h. To quantify the total biomass, the pegs with biofilms were stained using 0.1 % crystal violet (CV) (Sigma-Aldrich), and the absorbance (540 nm) of the dissolved stain was measured as described before^38^.

### Bacterial survival in human serum

O/N cultures were grown in LB and diluted 100-fold in PBS. A final dilution of ∼10^5^ CFU was mixed with human serum (Sigma-Aldrich, H6914 from male AB clotted whole blood, USA origin, sterile-filtered) in a total volume of 200 μl in a 96-well plate. Due to the natural batch variation of human serum, serum was diluted up to 40 % when needed to achieve a similar killing effect relative to the index and sensitive control isolates. For negative controls, serum was heat-inactivated for 30 min at 56°C. To confirm that the killing effect was due to the complement system, a control with the complement inhibitor polyanetholesulfonic acid (Sigma-Aldrich) was done at a final concentration of 100 µg/ml (Supplementary Fig. 5). CFU counts were performed at timepoint 0h and after 3h static incubation at 37°C.

### Virulence testing in *Galleria mellonella* larvae model

Research-grade wax moth *G. mellonella* larvae TruLarv^TM^ were purchased from BioSystems Technology Ltd (UK). O/N cultures (BHI) were diluted in PBS, and 10 μl containing 10^4^ CFU were injected via the first right proleg using a 25 µl Hamilton 7000 syringe (Model 702 RN, 22S gauge needle). A dose of 10^4^ CFU was chosen because it led to the most consistent results between biological replicates and the best resolution between different virulence levels. For each strain, two biological replicates (cultures started from independent colonies) were injected into 10 larvae each. For each experiment, 10 larvae were injected with 10 μl sterile PBS to account for possible injection trauma, and 10 larvae were used as non-injected controls. Aliquots from dilutions were plated for CFU counts to confirm the correct infection load. Larvae were incubated at 37°C for 48 h with an hourly assessment of death during 7-12 h post-injection, followed by monitoring after 24 h, 32 h, and 48 h. Larvae were considered dead when not moving after stimulation with a sterile pipette tip, which usually coincided with melanization. At the end of the experiment, live larvae were given health index scores^79^, and a cumulative score (0-90) was determined by adding up scores of individual larvae for each replicate. Survival analysis was performed using the Kaplan-Meier method and analyzed by log-rank (Mantel-Cox) test in GraphPad Prism version 9.2.0 for macOS (GraphPad Software, San Diego, CA, USA). All isolates were categorized relative to the index isolate DA14734.

### Construction of a genetically marked index isolate for mice experiments

The index isolate DA14734 was transformed with the pSIM10-hygro λ Red recombineering system. The aminoglycoside resistance gene *armA* was PCR amplified using Phusion High-Fidelity DNA Polymerase (Thermo Fisher Scientific Inc.) with primers introducing a 40 bp homology region directly outside *galK* for in-frame replacement (primers: 14734_armA_galK_F1 GCGCCGTCAGCGACGTCCATTTTCGTGAATCCGGAGTGTAAGTAAAGTGTGGTGATG TTGA;14734_armA_galK_R1_ATCGTCCGGGGCCAGCGCGGTGGTTTGCGTTAGCATTGTTGTCGTTTGTGAGCT. Purified *armA(galK)* DNA was electroporated into electrocompetent DA14734 pSIM5-hygro λ Red cells washed with 10% glycerol that had the λ Red system induced by 15 min incubation at 42°C. Successful transformants carrying *galK*::*armA* were selected on LA + gentamicin (50 mg/L) at 30°C.

### Competitions during gastrointestinal colonization in mice

To assess the competitive advantage during gastrointestinal colonization in mice, the marked index isolate DA73939 (*galK::armA*) was competed with the unmarked selected isolates. Testing all isolates in mice was not feasible, therefore, we chose five isolates DA14742, DA25061, DA25066, DA25071, and DA25116, representing different isolation sites, capsule status, biofilm formation capacity, virulence in *G. mellonella* larvae, and survival in human serum. BALB/c female mice (n=36) were acclimatized for eight days before starting the experiment. One mouse injured itself during this period and had to be sacrificed by cervical dislocation. As a result, one competition group (control DA14734 vs DA73939) had five instead of six mice. Overnight cultures of pair-wise isolates were mixed 1:1 in PBS/1% bicarbonate, and 100 µl of the mixture (approximately 3×10^7^ CFU) were administered via oral gavage. The procedure was repeated on two consecutive days (at least 24 h apart). On Day 5, a third gavage with fresh bacterial cultures was given. After that, drinking water with clindamycin (Villerton) at a final concentration of 0.5 g/L and 5 g/L glucose was provided *ad libitum* to disrupt the endogenous microflora and facilitate colonization. Feces from each mouse were collected every day (Day 1-Day 10), weighed, homogenized in 1 ml sterile PBS, vortexed at full speed for 2 min, and briefly (max 5 s) spun down to remove larger debris. Different dilutions were plated on LB agar with cefotaxime (10 mg/L) to allow growth of all isolates and LB agar with gentamicin (50 mg/L) to select for the marked index isolate. The number of bacteria was expressed as CFU per gram of feces. On Day 10, all mice were sacrificed by cervical dislocation. To test for systemic spread of bacteria, spleens were extracted, homogenized in 1 ml PBS, vortexed at full speed for 2 min, briefly spun (5 s), and plated on LB agar with cefotaxime (10 mg/L) or gentamicin (50 mg/L). Colonized mice showed no signs of disease or systemic infection at the end of the competitions. The competitive index was calculated each day by dividing the output ratio in feces by the input ratio (inoculum cultures) for each mouse. The final data was normalized to the values from the control competition to account for any possible selective differences conferred by *galK*::*armA* and expressed as a log_10_ competitive index. All experiments with the animals were done in accordance with the national regulations and approved by the Regional Ethics Review Board in Uppsala (Dnr 5.8.18-15552/2019).

### Growth under iron-depleted and repleted conditions

Iron-dependent growth in all isolates was assessed similarly as described before^34^. The iron-depleted medium was made by treating M9 with Chelex 100 (50-100 dry mesh) resin (BioRad) for 1 h, then filter-sterilized and supplemented with 0.2% glucose, 1 mM MgSO_4_ and 0.3 mM CaCl_2._ Overnight cultures were grown in BHI and diluted to 10^3^ CFU/ml in M9/Chelex, M9/Chelex supplemented with 100 µM FeSO_4,_ and M9/Chelex with 5% heat-inactivated human serum. The growth was monitored in a BioScreen C (Oy Growth Curves Ad Ltd) for 18 h at 37°C with shaking and OD_600_ measurements taken every 4 min. The index isolate was monitored along with the experimental isolates for each run. The area under the curve (AUC) was calculated for each isolate and each replicate, and then normalized to the index for each condition. AUC of each isolate was compared to the index isolate in the respective conditions by Brown-Forsythe and Welch ANOVA followed by Dunnett’s T3 multiple comparison test.

### RNA extraction and qPCR of *fepA* and *fes*

Overnight cultures of selected isolates were grown in BHI and diluted to 10^4^ CFU/ml in M9/Chelex, M9/Chelex supplemented with 100 µM FeSO_4,_ and M9/Chelex with 5% heat-inactivated human serum. The cultures were grown until the mid-exponential phase, spun down, and mixed with RNA Protect (Qiagen). RNA was extracted using the RNeasy Mini Kit (Qiagen), and genomic DNA was removed using the TURBO DNA-free kit (Invitrogen). Complementary DNA (cDNA) was synthesized using the High-Capacity Reverse Transcriptase kit (Applied biosystems) with approximately 500 ng of extracted RNA. The mRNA levels of *fepA* and *fes* were quantified by qPCR (Illumina Eco Real-Time PCR System), using the SYBR Green Master mix (Thermo Fisher) with the following primers: fepA_F1 CAGAACACCAACTCCAACGA; fepA_R1 CCGTTCCAGGTAACCGAGTA; fes_F1 AATCGCTGCAACGTCTCC, fes_R2 TCACGGTCGGAGGGAATA. The mRNA levels of the housekeeping gene *glnA* were assessed using primers glnA_F and glnA_R^80^. The mRNA levels were normalized to the index mRNA levels for the respective conditions using the 2^−ΔΔ*C*^ method^81^.

### Sedimentation assay

Hypermucoviscosity was assessed by low-speed centrifugation as described before^64^. Overnight cultures grown in BHI (5 ml, in 50 ml Falcon tubes) were centrifuged for 5 min at 1000x g, and OD_600_ values were measured, before and after centrifugation, for the top 300 µl of supernatants. Hypermucoviscosity was expressed as the ratio between the OD_600_ of the supernatant and the OD_600_ of the corresponding culture before centrifugation.

### Statistical analysis

All statistical analysis was done using GraphPad Prism version 9.2.0. for macOS (GraphPad Software, San Diego, California, USA). The respective analysis is specified for each experiment and figure in the figure legends.

## Acknowledgments

We thank Evelina Kess for handling the work with mice during the colonization experiments, and the staff at the clinical microbiology lab at Uppsala University Hospital for help with collection of clinical isolates. We also thank Dan I. Andersson and Vaughn S. Cooper for comments on the manuscript draft, and Lionel Guy for help with the phylogenetic analysis.

## Author contributions

G.Z. and L.S. conceived the study and designed experiments. G.Z. performed all phenotypic characterization experiments. K.H. performed whole-genome sequencing. G.Z., K.H., and L.S. performed bioinformatic analysis. B.L. provided clinical isolates and patient data. G.Z. and L.S. analyzed data and wrote the manuscript. All authors read, edited, and approved the final version of the manuscript.

