## Supplementary figures for "Convergent within-host evolution alters key virulence factors in a *Klebsiella pneumoniae* clone during a large hospital outbreak"

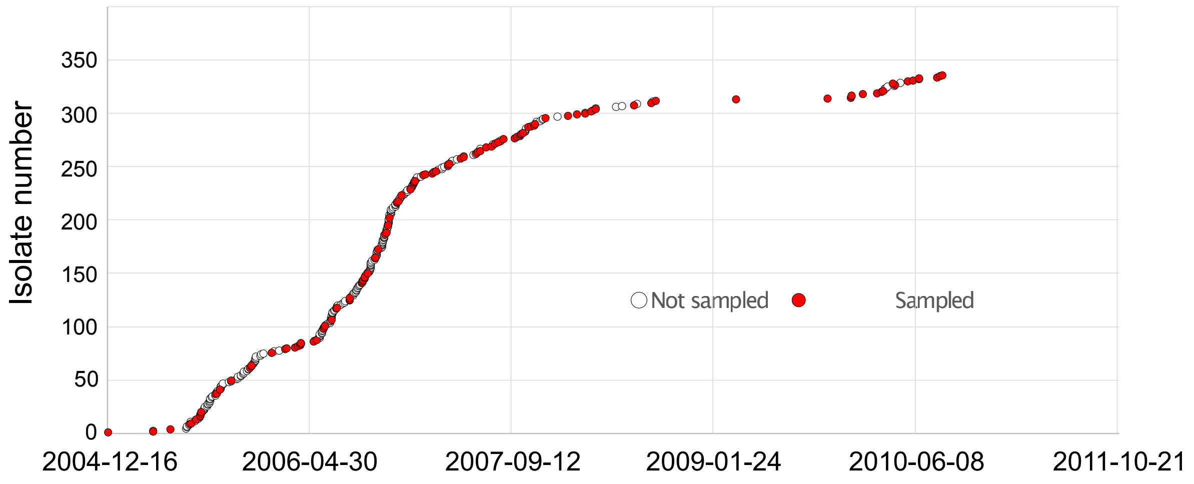


**Supplementary Fig. 1.** Cumulative sampling graph of isolates over time of the outbreak. Red circles denote sampled isolates in this study.

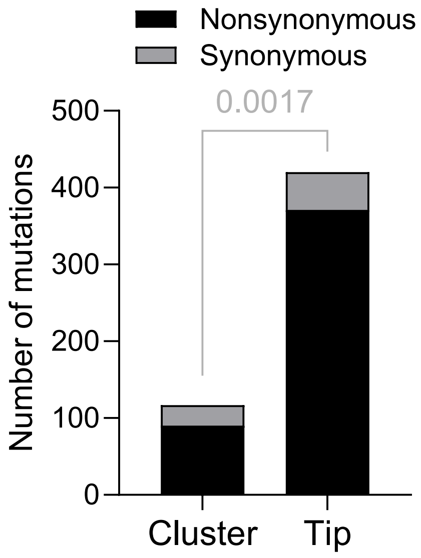


**Supplementary Fig. 2.** Distribution of nonsynonymous and synonymous mutations in cluster and tip mutations. Siginficance assessed by Chi-square, p value shown above the bars.

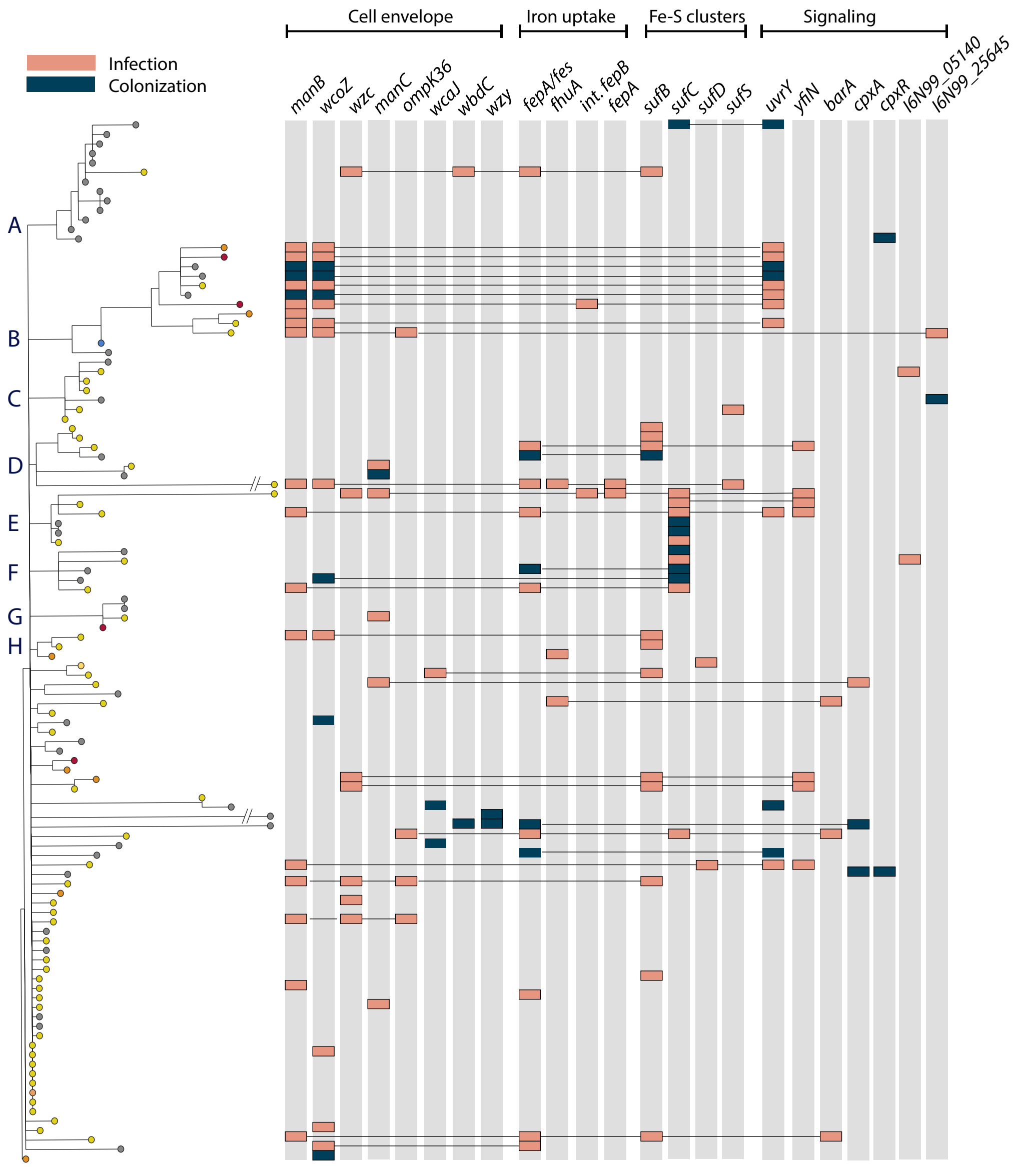


**Supplementary Fig. 3.** The most commonly mutated gene-combinations on the phylogenetic tree. Letters denote clusters corresponding to Fig. 1c. Full figure also available in Supplementary Table 3.

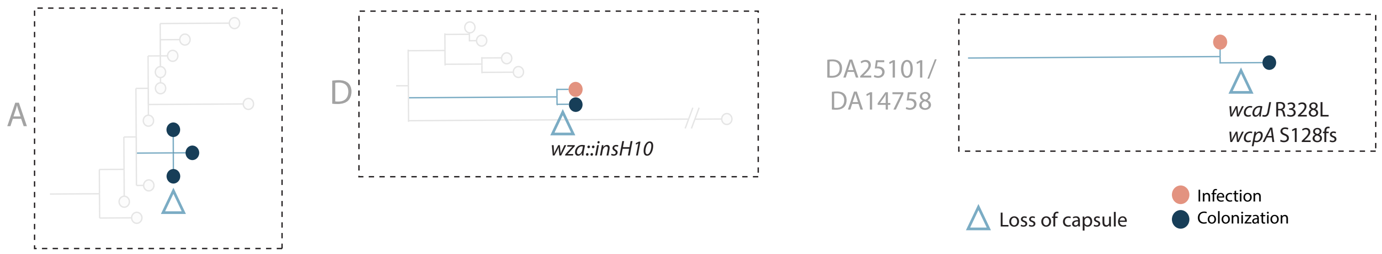


**Supplementary Fig. 4.** Phylogenetic clusters A, D, and DA25101/DA14758 with attenuated/slower killing isolates where the capsule was lost at a later timepoint.

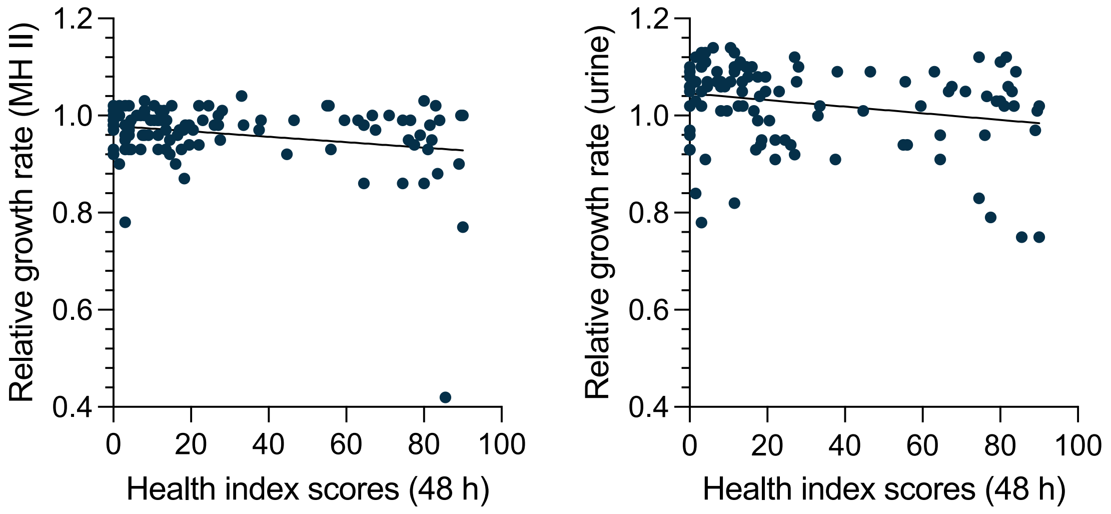


**Supplementary Fig. 5.** Exponential growth rates in MH II medium (left) and urine (right) relative to the index isolate vs health index scores in *G. mellonella* larvae at 48 h post-injection.

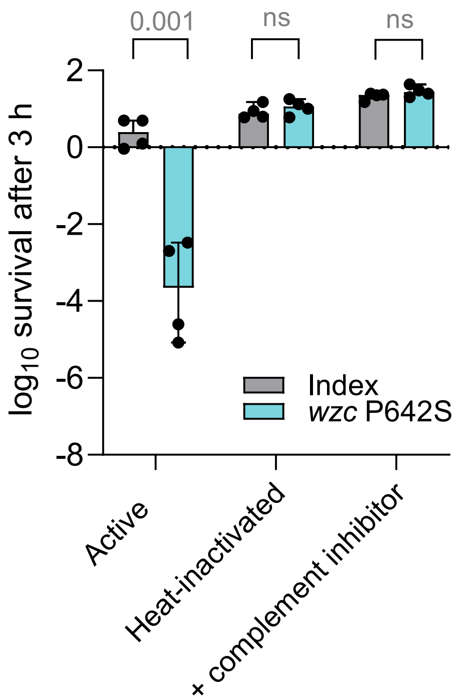


**Supplementary Fig. 6.** Serum killing control with active serum, heat-inactivated serum and complement inhibitor polyanetholesulfonic acid. Index refers to the index isolate, *wzc* P642S - isolate with a single mutation in *wzc*. Results from four biological replicates with 95% CI are shown. Difference between the index isolate index and *wzc* P642S mutant in each condition tested by unpaired t-test, ns - not significant.
